# The interplay between CB_2_ and NMDA receptors in Parkinson’s disease

**DOI:** 10.1101/2025.01.20.633878

**Authors:** Jaume Lillo, Iu Raïch, Joan Biel Rebassa, Toni Capó, Pau Badia, Gemma Navarro, Irene Reyes-Resina

**Author notes:** Correspondence (I.R.-R.); (G.N.).

## Abstract

Parkinson’s disease (PD) is a progressive neurological disorder that affects movement, causing symptoms such as tremors, stiffness, slowness, and balance problems due to the degeneration of dopamine-producing neurons in the brain. Nowadays there is no cure for PD. Alpha synuclein (α-syn) aggregates, which are a hallmark of PD, are known to induce microglial activation, specifically the detrimental M1 microglial phenotype, which contributes to neuroinflammation and disease progression. Cannabinoid receptor 2 (CB_2_R) activation has been shown to counteract neuroinflammation. CB2R is able to interact with NMDA receptors (NMDAR), which has also attracted attention in PD research due to its role in excitotoxicity. Here we aimed to study the interaction between CB_2_R and NMDAR in a PD context. We observed that α-syn fibrils alter CB2R activation and CB_2_R-NMDAR heteromerization in a heterologous expression system. Furthermore, activation of CB2R counteracted NMDAR signaling. In neurons α-syn fibrils decreased CB2R-NMDAR heteromer expression, while increasing CB_2_R signaling. Importantly, CB_2_R activation counteracted the α-syn fibrils-induced increase in M1 activated microglia, while it favored the polarization of microglia to the beneficial M2 phenotype. These results reinforce the idea of using cannabinoids for treating PD, as they provide not only the anti-inflammatory effects of cannabinoids but also counteract the detrimental increase in NMDAR signaling present in this disease.

## 1. INTRODUCTION

Parkinson’s disease is the most common neurodegenerative movement disorder (de Rijk et al., 1997). In Europe, prevalence and incidence rates for PD are estimated at approximately 200/100 000 and 15/100 000 per year, respectively (Balestrino & Schapira, 2020). Age and male gender are considered risk factors (Van Den Eeden, 2003). This pathology curses with the loss of the dopaminergic neurons of the substantia nigra pars compacta, whose axons project to the striatum. As a consequence of the lack of dopamine in the striatum the direct and indirect pathways are dysregulated, symptoms such as bradykinesia, rigidity, tremor and postural instability appear (Poewe et al., 2017). The etiology of the disease in most patients is unknown. The familial form of Parkinson’s disease accounts for only 14 % of all cases, and it has been linked to mutations in different genes, such as alpha-synuclein (α-syn), parkin, DJ-1, PINK1, and LRRK2 (Klein & Westenberger, 2012). The majority of PD cases are sporadic, being associated with variants of genes that can influence the susceptibility to PD, which have been suggested to be caused by interaction with environmental toxins (Ascherio & Schwarzschild, 2016).

PD is characterized by the presence of Lewy bodies, which are intracellular inclusions present in the remaining dopaminergic neurons, composed of aggregates of misfolded α-synuclein (Calabresi et al., 2023). This presynaptic protein seems to have a role in the biosynthesis and liberation of neurotransmitter vesicles, by promoting SNARE-complex assembly during vesicle docking and fusion steps (Yoo et al., 2023). In both PD forms the presence of α-syn aggregates has been described. The misfolded form of α-syn, that appears due to mutations or polymorphisms in the SNCA gene (Polymeropoulos et al., 1997; Simón-Sánchez et al., 2009), aggregates forming oligomers, which will further aggregate in the form of fibrils, that finally generate Lewy bodies (Spillantini et al., 1997). Current treatment for PD is based on the replacement of dopamine, being the most common treatment the dopamine precursor levodopa, which produces secondary effects such as the abnormal involuntary movements known as dyskinesias (Ahlskog & Muenter, 2001). Alternative approaches such as deep brain stimulation (DBS) are suitable for later-stage disease. Current available treatments offer good control of motor symptoms but do not halt the progression of neurodegeneration, the evolution of the disease and the increasing disability.

Cumulative evidence points to a role of α-synuclein in activating immunological response, as it has been described that Lewy body-like alpha-synuclein inclusions trigger reactive microgliosis prior to nigral degeneration (Duffy et al., 2018). Furthermore, it has been demonstrated that activated microglial cells directly engulf α-synuclein in an attempt to clear it (Rocha et al., 2018). Indeed, it is known that neurodegenerative diseases such as PD or AD course with neuroinflammation. Postmortem analyses in PD patients revealed the presence of activated microglia and increased expression of inflammatory markers in the substantia nigra, putamen and cortex (Imamura et al., 2003; McGeer et al., 1988). Analysis of PD patients post-mortem brain tissue revealed a between microglial activation and α-synuclein pathology in the SN and the hippocampus (Imamura et al., 2003). A recent study proposes a model in which α-syn accumulation activates the detrimental M1 type microglia, while it does not activate the M2 type, which is considered neuroprotective (Su & Zhou, 2021).

Cannabinoids have been proposed to decrease neuroinflammation in neurodegenerative diseases (Chiurchiù et al., 2015; Pérez-Olives et al., 2021). Cannabinoids mainly act on cannabinoid CB_1_R and CB_2_R receptors, being CB_2_R of great interest, as its activation lacks psychoactive effects (Lu & Mackie, 2021). CB_2_R expression is higher in glial cells than in neurons, and it increases upon microglial activation, which seems to underlie the neuroprotective effects of cannabinoids (D. Chen et al., 2017; Navarro et al., 2018). Furthermore, an increase of CB_2_R expression was found in microglial cells in the post-mortem substantia nigra of PD patients and in the striatum and the substantia nigra in lipopolysaccharide (LPS)-lesioned mice (Gómez-Gálvez et al., 2016). The 1-methyl-4-phenyl-1,2,3,6-tetrahydropyridine (MPTP) neurotoxic mouse PD model shows a significant microglial activation and overexpression of CB_2_ receptors in midbrain (Price et al., 2009). This increase in CB_2_R expression was also found in the neurotoxic 6-hydroxydopamine (6-OHDA), the environmental and the inflammation-driven rat models of Parkinson’s disease, and this change significantly correlated with an increase in microglial activation (Concannon et al., 2015, 2016). The pharmacological activation of CB_2_ receptors has been shown to reduce proinflammatory responses in the LPS-induced mouse model (Gómez-Gálvez et al., 2016), in the MPTP neurotoxic mouse models (Price et al., 2009) and in the rotenone rat model (Javed et al., 2016) of PD, and thus to protect dopaminergic neurons from degeneration. On the contrary, the genetic inactivation of CB_2_ receptors aggravated LPS-induced inflammation (Gómez-Gálvez et al., 2016) and led to a more pronounced MPTP-induced toxicity in the substantia nigra (Price et al., 2009), respectively. Hence, this receptor has been proposed as therapeutic target for Parkinson’s disease (Abellanas & Aymerich, 2020; Gómez-Gálvez et al., 2016).

The ability of GPCRS to form complexes with other membrane receptors is well known (Borroto-Escuela et al., 2017). Cannabinoid receptors CB_1_ and CB_2_ have been described to form complexes, among others, with N-methyl-D-aspartate receptors (NMDARs) in the central nervous system, both in neurons and in glia (Reyes-Resina et al., 2024; Rivas-Santisteban et al., 2021). CB_1_R-NMDAR complexes seem to play an important role in Parkinson’s (Reyes-Resina et al., 2024) disease, however, the role of CB_2_R-NMDAR complexes in a Parkinson’s disease context remains unexplored (Rivas-Santisteban et al., 2021).

Furthermore, the alteration in the expression and function of NMDARs is other of the hallmarks of PD. NMDARs are heteromeric glutamate-gated ion channels formed by four subunits, which in humans are usually two obligatory NR1 subunits plus two NR2A or NR2B subunits (Paoletti et al., 2013). They are implicated in synaptic plasticity, learning, and memory but also in neurodegeneration and excitotoxicity (Bading, 2017; Hardingham & Bading, 2010; Parsons & Raymond, 2014). In the rat PD model consisting of unilateral nigrostriatal dopamine system ablation with 6-OHDA, the NMDAR phosphorylation state is altered (Dunah et al., 2000; Oh et al., 1998). Also, in PD patients and in animal models of the disease, an alteration in the expression levels of NMDAR subunits has been found (Dunah et al., 2000; Gan et al., 2014; Mellone et al., 2015). Interestingly, NMDAR antagonists such as MK-801, or negative allosteric modulators such as radiprodil, have been shown to improve motor symptoms in PD (Melo-Thomas et al., 2018; Michel et al., 2017). However, there is currently no cure for Parkinson’s disease, only symptomatic treatments are available, often with undesired secondary effects (Armstrong & Okun, 2020).

Given the lack of a cure for Parkinson’s disease, our aim was to determine if CB_2_R-NMDAR complexes expressed in microglia could have a role as therapeutic targets in PD. We studied the expression and function of these complexes in a heterologous expression system and in an in vitro model of PD consisting of primary cultures of microglia treated with α-syn.

## 2. RESULTS

### 2.1. α-syn fibrils alter the conformation of CB_2_R-NMDAR complexes

To determine if CB_2_R-NMDAR complexes are altered in a parkinsonian context, we first studied if preformed α-syn fibrils could alter the localization of CB_2_R and NMDAR in the plasma membrane. HEK-293T cells have been previously described be able to uptake these α-syn preformed fibrils after a 2-day treatment (Reyes-Resina et al., 2024). HEK-293T cells expressing either CB_2_R fused to the fluorescent protein YFP, NMDAR fused to the Renilla luciferase (Rluc), or both, were analyzed under the confocal microscope. CB_2_R expression was detected by the YFP’s own fluorescence, while expression of NMDAR was detected by immunocytochemistry with an anti-Rluc antibody plus a Cy3 conjugated secondary antibody. The expression of both receptors was shown at the plasma membrane level of HEK-293T cells, and when cells were co-expressing both receptors, colocalization was observed (Fig. 1A). When the same experiment was performed in the presence of α-syn fibrils, no differences were found compared to the control condition, neither in the expression nor in the colocalization between CB_2_R and NMDAR (Fig. 1A). To rule out the possibility that α-syn fibrils cause HEK-293T cell death, cells were incubated with increasing concentrations of α-syn fibrils for 48 h. When HEK-293T cell viability was measured, we observed that no cell death was caused by α-syn fibrils (Fig. 1B).

**Figure 1.**
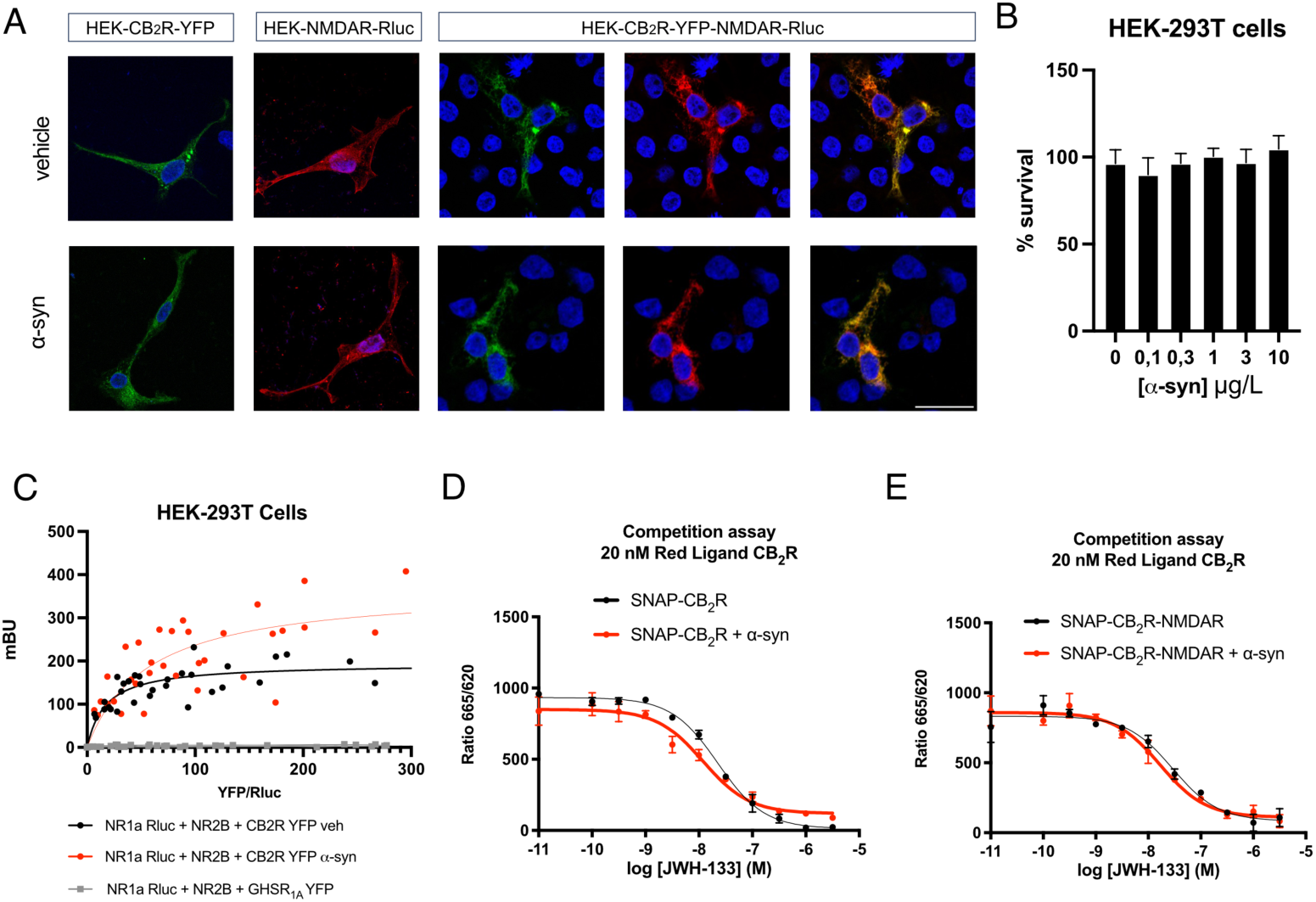
Analysis of CB_2_R-NMDAR complex formation in the presence of α-syn. **(A)** Immunocytochemistry assay was performed in HEK-293T cells treated or not with 10 µg/L of human α-syn fibrils for 48 h and transfected with either CB_2_R-YFP (1 µg cDNA) (shown in green), NR1-Rluc (0,75 µg cDNA) (shown in red) plus NR2B (0.75 µg cDNA), or both. Nuclei were stained with Hoechst (blue). Colocalization is shown in yellow. Scale bar: 20 µm. **(B)** HEK-293T cells were treated with increasing concentrations of α-syn or vehicle for 48 h. Then, cells were gently detached, and cell survival was assessed with a cell counter. Values are the mean ± SEM of 5 independent experiments performed in triplicates. One-way ANOVA, followed by Tukey’s multiple comparison post hoc test, was used for statistical analysis. **(C)** BRET assays were performed in HEK-293T cells treated or not with 10 µg/L of human α-syn fibrils for 48 h and transfected with constant amounts of cDNAs for NR1-Rluc (0.5 µg) and NR2B (0.3 µg) and increasing amounts of cDNA for CB_2_R-YFP (0 to 2.5 µg) or GHSR_1A_-YFP (0 to 10 µg). Values are the mean ± SEM of 5 different experiments performed in triplicates. **(D, E)** HTRF-based competitive binding assays were performed in HEK-293T cells transiently transfected with 1.5 µg cDNA for SNAP-CB_2_R in the absence **(D)** or presence of 0,7 µg cDNA for NR1 and 0,7 µg cDNA for NR2B **(E)**. Competition curves of specific binding of 20 nM fluorophore-conjugated CM157 using JWH-133 as competitor are shown. Data represent the mean ± SEM of five experiments performed in triplicates.

Next, we aimed to study if α-syn fibrils could alter the formation of CB_2_R-NMDAR complexes. Thus, the Bioluminiscence Resonance Energy Transfer (BRET) technique was carried out to assay physical interactions between receptors in vivo. When we performed a BRET assay in HEK-293T cells expressing constant amounts of NMDAR fused to Rluc and increasing amounts of CB_2_R fused to YFP, a saturation curve was obtained (Fig. 1C), which indicates that CB_2_R and NMDAR can physically interact, as previously described (Rivas-Santisteban et al., 2021). The obtained BRET_max_ was 200 ± 27 mBU (milli BRET units), and the BRET_50_ was 17 ± 10. However, when the same assay was performed in the presence of α-syn fibrils, a higher BRET signal was observed (Fig. 1C), with a BRET_max_ of 370 ± 90 mBU (milli BRET units) and a BRET_50_ of 56 ± 40. These results demonstrate that α-syn fibrils do not disrupt the formation of CB_2_R-NMDAR complexes. Instead, fibrils likely cause a protein reorganization at the plasma membrane that changes the orientation of the Rluc and YFP, resulting in a better energy transfer between them, as described for CB_1_R-NMDAR complexes (Reyes-Resina et al., 2024). To show that the interaction between CB_2_R and NMDAR is specific, a negative control experiment was performed, in which CB_2_R-YFP was substituted by the ghrelin receptor GHSR_1A_ fused to YFP, which resulted in a linear signal (Fig. 1C), indicative of a lack of interaction between NMDAR and GHSR_1A_.

### 2.2. α-syn fibrils do not affect CB_2_R affinity for its ligands

A common phenomenon in GPCR biology is the fact that the interaction with other receptors affects the pharmacology of GPCRs. Thus, we first aimed to determine if the interaction of CB_2_R with NMDAR could alter ligand binding to the orthosteric site of CB_2_R. Homogeneous Time-Resolved FRET (HTRF) assays were performed in HEK293-T cells expressing either only CB_2_R fused to the SNAP protein (SNAP-CB_2_R) or SNAP-CB_2_R together with NMDAR, where increasing amounts of the selective CB_2_R agonist JWH-133 (from 0,1 nM to 10 μM) were used to displace the fluorophore-conjugated selective CB_2_R agonist CM157. In both cell types JWH-133 decreased the binding of labeled CM157 to SNAP-CB2R in monophasic fashion (Fig. 1D, E). The obtained Ki values were 18 and 23,4 nM, respectively, indicating that the interaction with NMDAR does not significantly alter the affinity of CB_2_R for JWH-133. Next, we sought to determine how α-syn fibrils affected the binding of ligands to CB_2_R. When HTRF assays were performed after incubation with fibrils, again monophasic competition curves were obtained (Fig. 1D, E), with Ki values of 9,63 nM for cells expressing SNAP-CB_2_R and 13,5 nM for cells expressing SNAP-CB_2_R plus NMDAR. These results indicate that α-syn fibrils does not affect the affinity of CB_2_R for its selective agonist JWH-133.

### 2.3. CB_2_R signaling is decreased upon treatment with α-syn fibrils

After studying how α-syn fibrils affect the formation of CB_2_R-NMDAR heteromers in HEK-293T cells, we wondered how fibrils could affect the functionality of these receptor complexes. Firstly, we determined the effect α-syn fibrils in the signaling of individual CB_2_R and NMDAR. As CB_2_R is a GPCR coupled to Gi protein, intracellular cAMP levels were assayed. In HEK-293T cells expressing CB_2_R, the selective CB_2_R agonist JWH-133 was able to induce a 50% decrease in intracellular cAMP levels previously increased by pre-treatment with forskolin (Fig. 2A). When these cells were pre-treated with the CB_2_R antagonist SR144528, the effect of JWH133 was blocked (Fig 2A). However, when cAMP levels were tested after incubation with α-syn fibrils, the decrease in cAMP levels induced by JWH-133 was only a 30% (Fig. 2B). The recruitment of β-arrestin II upon CB2R activation was also assayed. In HEK-293t cells expressing CB_2_R fused to YFP and β-arrestin II fused to Rluc a BRET assay was performed. CB_2_R agonist JWH-133 was able to induce a significant β-arrestin II recruitment, that was counteracted by CB_2_R antagonist SR144528 (Fig. 2C). The signal induced by JWH-133 was abolished upon α-syn fibrils treatment (Fig. 2D). These results indicate that α-syn fibrils reduce CB_2_R activation in HEK-293T cells, in line with the effects caused by α-syn fibrils on CB_1_R signaling (Reyes-Resina et al., 2024).

**Figure 2.**
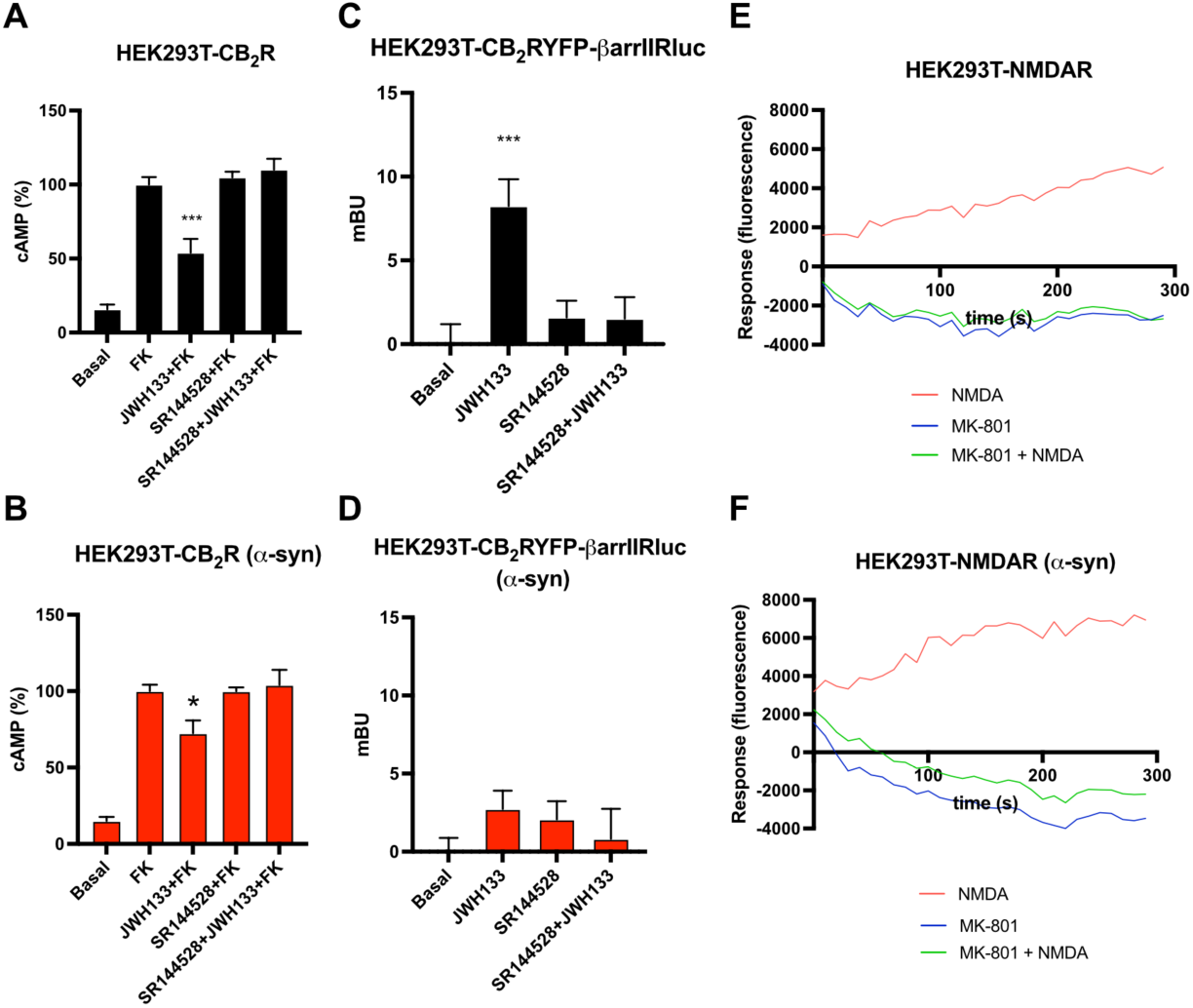
Analysis of CB_2_R and NMDAR signaling in the presence of α-syn fibrils in a heterologous system. **(A,B)** HEK-293T cells transfected with CB_2_R (1,3 µg cDNA) were treated **(B)** or not **(A)** with α-syn fibrils. Forty-eight hours after, cells were pre-treated with the vehicle or with the selective CB_2_R antagonists SR144528 1 µM, followed by agonist stimulation (100 nM JWH-133). cAMP accumulation was detected by HTRF in the presence of 0.5 µM forskolin. Values are the mean ± SEM of 6 different experiments performed in triplicates, and one-way ANOVA, followed by Tukey’s multiple comparison post hoc test, was used for statistical analysis (*p < 0.05, ***p < 0.001; versus treatment with forskolin). **(C, D)** HEK-293T cells transfected with CB_2_R-YFP (1,2 µg cDNA) together with β-arrestin II-Rluc (0,4 µg cDNA) were treated **(D)** or not **(C)** with α-syn fibrils. Forty-eight hours after, cells were pre-treated with the vehicle or with the selective CB_2_R antagonists SR144528 1 µM, followed by agonist stimulation (100 nM JWH-133). β-arrestin II recruitment was assayed by BRET assays. Values are the mean ± SEM of 6 different experiments performed in triplicates, and one-way ANOVA, followed by Tukey’s multiple comparison post hoc test, was used for statistical analysis (***p < 0.001; versus basal). **(E, F)** Calcium release was evaluated in HEK-293T transfected with NR1 (1 µg cDNA), NR2B (1 µg cDNA) and with 6GCamMP calcium sensor (0,3 µg cDNA) and treated **(F)** or not **(E)** with α-syn fibrils. Cells were pre-treated with vehicle or with the selective antagonists (1 µM MK-801), followed by agonist stimulation (15 µM NMDA). Data represent the mean ± SEM of six different experiments performed in triplicates.

Regarding NMDAR, ion channel activation has no effect neither in cAMP levels nor in β-arrestin II recruitment, hence intracellular Ca^2+^ mobilization was studied. When HEK-293T cells expressing NMDAR were treated with the NMDAR agonist, a calcium signal was observed, that was counteracted by pre-treatment with NMDAR antagonist MK-801 (Fig. 2E). When calcium mobilization was studied in the presence of α-syn fibrils, no difference was observed compared to cells treated with vehicle (Fig. 2F), indicating that α-syn fibrils do not affect intracellular Ca^2+^ mobilization upon NMDAR activation in HEK-293T cells.

### 2.4. Signaling of CB_2_R-NMDAR complexes is decreased upon α-syn fibrils treatment in a heterologous system

After studying the signaling on individual receptors, the effect of α-syn fibrils in the functionality of CB_2_R-NMDAR complexes was tested. Intracellular cAMP levels were assayed in HEK-293T cells expressing both CB_2_R and NMDAR. In these cells, the CB_2_R agonist JWH-133 induced a 40% decrease in cAMP levels previously increased by forskolin, while NMDAR agonist did not induce a significant effect (Fig. 3A). When cells were co-treated with both agonists together, a similar effect to that observed with JWH-133 alone was detected, indicating that NMDAR activation does not affect CB_2_R signaling (Fig. 3A). When cAMP levels were measured after incubation with α-syn fibrils, the signal induced by JWH-133 was similar to that obtained in the absence of fibrils, indicating that in CB_2_R-NMDAR complexes α-syn fibrils do not affect CB_2_R signaling in the intracellular cAMP pathway (Fig. 3B).

**Figure 3.**
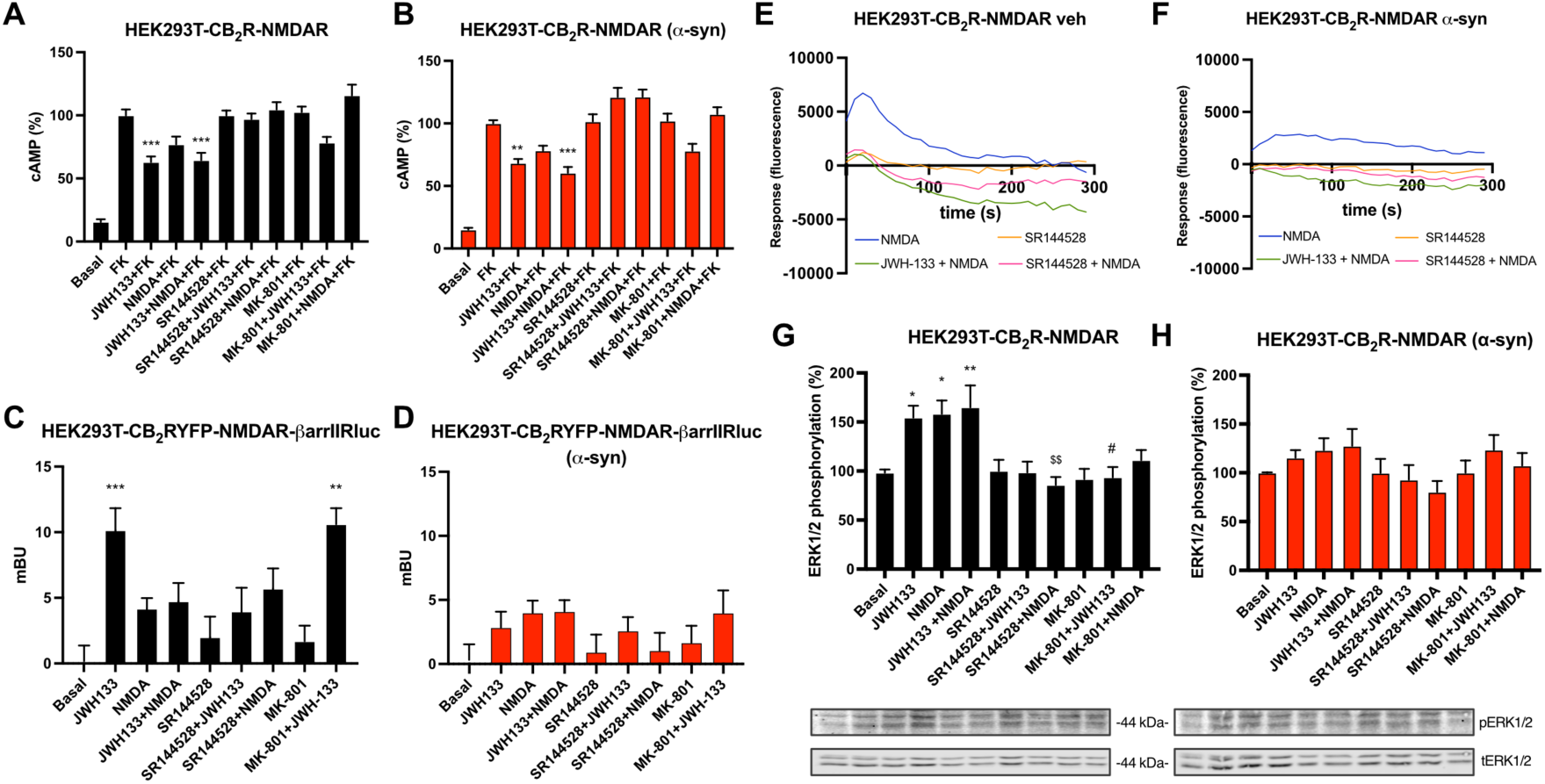
Functional analysis of CB_2_R-NMDAR complexes in a heterologous system upon α-syn treatment. **(A, B, G, H)** HEK-293T cells were transfected with the cDNAs for CB_2_R (1 µg), NR1 (0,7 µg), and NR2B (0,7 µg) and treated **(B, H)** or not **(A, G)** with α-syn fibrils. Cells were activated with the selective antagonists (1 µM SR144528 for CB_2_R or 1 µM MK-801 for NMDAR), followed by agonist stimulation (100 nM JWH-133 for CB2R and/or 15 µM NMDA for NMDAR). cAMP accumulation was detected by HTRF in the presence of 0,5 µM forskolin **(A, B)**. MAPK phosphorylation was detected by Western blot (blots for phosphorylated and total ERK1/2 (42–44 kDa) are shown below) **(G, H)**. Values are the mean ± SEM of 6 different experiments performed in triplicates, and one-way ANOVA, followed by Tukey’s multiple comparison post hoc test, was used for statistical analysis (*p < 0.05, **p < 0.01, ***p < 0.001; versus treatment with forskolin treatment in cAMP or versus basal in MAPK, # p < 0.05 versus JWH-133, $$ p < 0.01 versus NMDA). **(C, D)** HEK-293T cells transfected with CB_2_R-YFP (1,2 µg cDNA), NR1 (0,7 µg), and NR2B (0,7 µg) together with β-arrestin II-Rluc (0,4 µg cDNA) were treated **(D)** or not **(C)** with α-syn fibrils. Forty-eight hours after, cells were pre-treated the selective antagonists, followed by agonist stimulation. β-arrestin II recruitment was assayed by BRET assays. Values are the mean ± SEM of 6 different experiments performed in triplicates, and one-way ANOVA, followed by Tukey’s multiple comparison post hoc test, was used for statistical analysis (**p < 0.01, ***p < 0.001; versus basal). **(E, F)** Calcium release was evaluated in HEK-293T cells transfected with the cDNAs for CB_2_R (1 µg), NR1 (0,7 µg) NR2B (0,7 µg), and 6GCamMP calcium sensor (0,3 µg) and treated **(F)** or not **(E)** with α-syn fibrils. Cells were activated with the selective antagonists, followed by agonist stimulation. Data represent the mean ± SEM of six different experiments performed in triplicates.

β-arrestin II recruitment was studied in HEK-293T cells expressing CB_2_R fused to YFP together with NMDAR, and β-arrestin II fused to Rluc. Stimulation with CB_2_R agonist JWH-133 yielded a significant signal, while, as expected, NMDA treatment induced no response (Fig. 3C). Co-treatment with both JWH-133 and NMDAR abolished JWH-133-induced signal (Fig. 3C), indicating that NMDAR activation decreases recruitment of β-arrestin II by CB_2_R. Upon α-syn fibrils treatment, JWH-133 was not able to induce a significant β-arrestin recruitment (Fig. 3D), indicating that in this pathway α-syn fibrils disrupt CB_2_R signaling.

To gain insights in NMDAR signaling, intracellular Ca^2+^ mobilization assays were performed. HEK-293T cells expressing CB_2_R, NMDAR and a calcium sensor protein (T.-W. Chen et al., 2013) responded to NMDA stimulation, and this signal that was significantly decreased by co-treatment with JWH-133, which indicates that CB_2_R activation decreases NMDAR signal (Fig. 3E). NMDA-induced signal was counteracted by CB_2_R antagonist SR144528 (Fig. 3E), pointing to a cross-antagonism effect from CB_2_R to NMDAR in CB_2_R-NMDAR complexes in HEK-293T cells. Treatment with α-syn fibrils induced a decrease in NMDAR signaling (Fig. 3F).

A signaling pathway involved in both GPCR and NMDAR activation is the MAP kinase (MAPK) phosphorylation pathway. When HEK-293T cells expressing CB_2_ and NMDAR were stimulated with either the selective CB_2_R agonist JWH-133 or with NMDA, a significant increase in ERK1/2 phosphorylation was detected (Fig. 3G). When cells were treated with both agonists together, a non-additive effect was observed (Fig. 3G). Pre-treatment with CB_2_R antagonist SR144528 and with NMDAR antagonist MK-801 blocked both JWH-133 and NMDA-induced signals (Fig. 3G), indicating that there is a bidirectional cross-antagonism phenomenon in MAPK signaling pathway. When ERK1/2 phosphorylation was tested in the presence of α-syn fibrils, no significant signals were obtained with any of the agonists (Fig. 3H), thus fibril treatment abolishes both CB_2_R and NMDAR-mediated MAPK activation in HEK-293T cells.

### 2.5. The expression and function of CB_2_R-NMDAR complexes are altered by α-syn fibrils in microglia

To study how α-syn fibrils affect CB_2_R-NMDAR complexes in a more physiological context, we next studied the effects of α-syn fibrils on these heteromers in microglial cells, as they are key players in the inflammation process, where CB_2_R has an important role, as mentioned above. Therefore, rat microglia primary cultures were treated with α-syn fibrils. Immunocytochemistry with an anti-human alpha synuclein antibody was used to demonstrate that microglial cells are able to uptake α-syn fibrils from the extracellular space, as a high fluorescent signal was detected in cells treated with fibrils compared to vehicle-treated cells (Fig. 4A). Cell survival assays showed that α-syn fibrils do not significantly affect the viability of microglia (Fig. 4B). Treatment with CB2R agonist ACEA did not induce any change in cell survival, while treatment with NMDA decreased cell viability (Fig. 4B).

**Figure 4.**
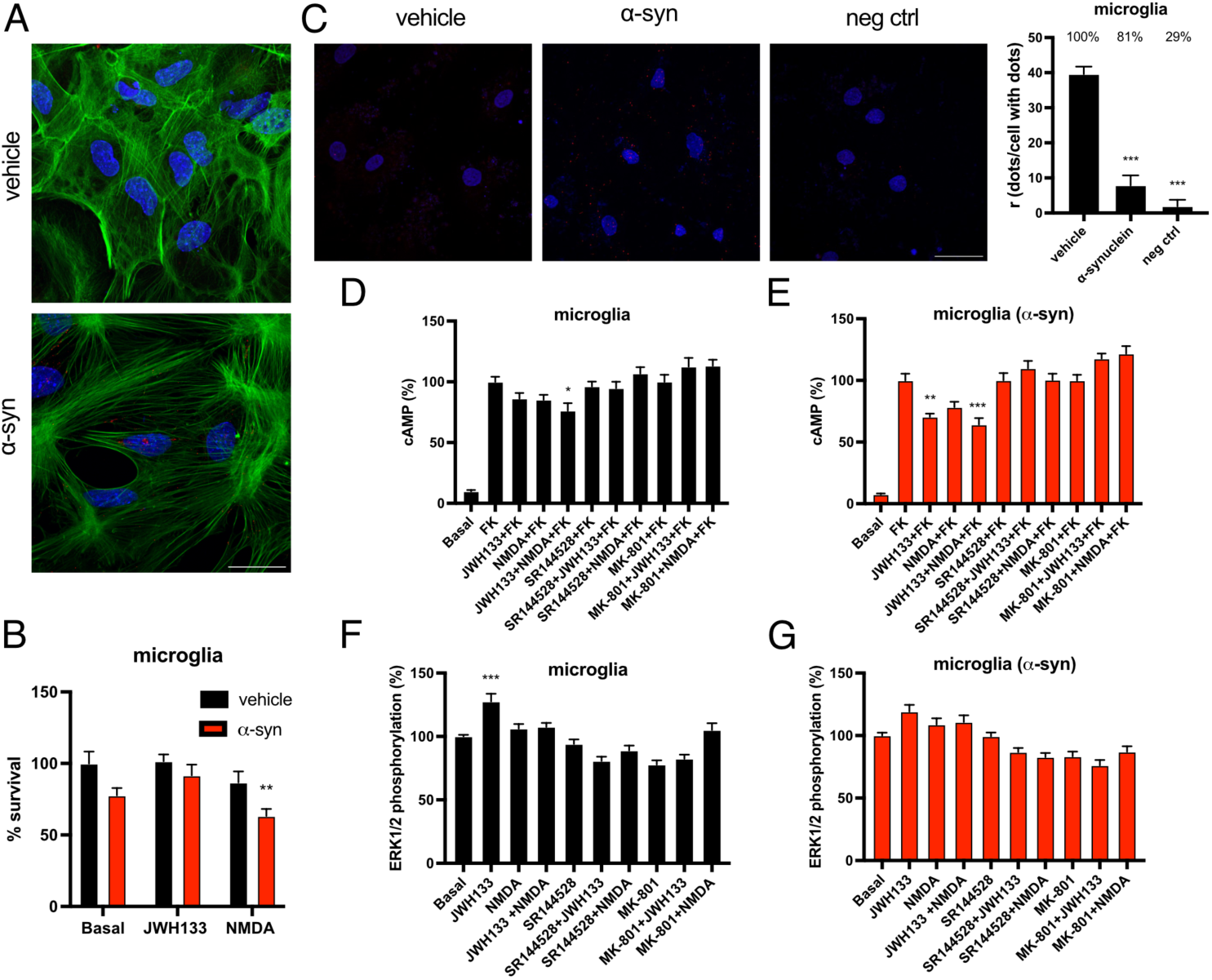
Expression and functionality of CB_2_R-NMDAR complexes in microglia treated with α-syn fibrils. **(A)** Primary cultures of microglia were treated with α-syn or vehicle. α-syn fibrils were detected by immunocytochemistry with a mouse anti-human α-synuclein anti-body (red). Nuclei were stained with Hoechst (blue). Scale bar: 30 µm. **(B)** Primary cultures of microglia were treated with α-syn or vehicle for 24 h, followed by treatment with JWH-133 (100 nM) or NMDA (15 mM) or vehicle for another 24 h period. Then, cells were gently detached, and cell survival was assessed with a cell counter. Values are the mean ± SEM of 5 independent experiments performed in triplicates. Two-way ANOVA, followed by Tukey’s multiple com-parison post hoc test, was used for statistical analysis (** p < 0.01, versus vehicle basal). **(C)** Proximity Ligation Assay (PLA) was performed in primary cultures of microglia treated or not with α-syn fibrils. Confocal microscopy images are shown (superimposed sections) in which heteromers appear as red clusters. Cell nuclei were stained with Hoechst (blue). Scale bar: 30 μm. Quantification of the number of red dots/cells with dots (r) and of the percentage of cells presenting red dots is shown. Values are the mean ± SEM (n = 6). One-way ANOVA, followed by Tukey’s multiple comparison post hoc test, was used for statistical analysis (*** p < 0.001, versus vehicle). **(D–G)** Primary cultures of microglia treated **(E,G)** or not **(D,F)** with α-syn were stimulated with the selective antagonists (1 µM SR144528 for CB_2_R or 1 µM MK-801 for NMDAR), followed by agonist stimulation (100 nM JEW-133 for CB_2_R and/or 15 µM NMDA for NMDAR) and cAMP levels **(D,E)**, and MAPK phosphorylation signals were measured **(F,G)**. Values are the mean ± SEM of 6 different experiments. One-way ANOVA, followed by Tukey’s multiple comparison post hoc test, were used for statistical analysis (* p < 0.05, **p < 0.01, *** p < 0.001; versus forskolin in cAMP assay or versus basal in MAPK phosphorylation).

To study heteromerization of CB_2_R and NMDAR in microglial cells, a proximity ligation assay (PLA) was performed in primary cultures of rat microglia. This technique allows the visualization of receptor-receptor complexes as red fluorescent dots under the microscope. Microglia cells showed around 40 dots/cell, with 100% of the cells showing fluorescent dots, while treatment with α-syn fibrils decreased PLA signal to 8 dots/cell with dots and only 80% of the cells presenting red dots (Fig. 4C). PLA negative control, in which one of the antibodies is omitted, resulted in 2 dots/cell with dots and only 30 % of the cells showing signal (Fig. 4C). Thus, α-syn fibrils treatment decreases the formation of complexes between CB_2_R and NMDAR.

Once we had determined how α-syn fibrils affected CB_2_R-NMDAR heteromerization, we aimed to address how these fibrils affect the functionality of the complex in microglia cells. When intracellular cAMP accumulation was analyzed, neither did CB_2_R nor NMDAR agonists produce a significant signal (Fig. 4D). However, upon α-syn fibrils treatment, a significant decrease in cAMP levels previously increased with forskolin was observed, both with CB_2_R agonist JWH-133 alone or in combination with NMDA (Fig. 4E), indicating that α-syn fibrils treatment increases CB_2_R signaling through the G-protein dependent pathway in microglial cells. Furthermore, JWH-133 signal was blocked not only by CB_2_R antagonist SR144528, but also by NMDAR antagonist MK-801 (Fig. 4D, E), indicating that there is a cross-antagonism effect from NMDAR to CB_2_R.

Next, we examined ERK1/2 phosphorylation in primary cultures of microglia by the alpha-screen method. When cells were stimulated with CB_2_R agonist JWH-133, a significant ERK1/2 was phosphorylation detected, while treatment with NMDA did not induce activation of this pathway (Fig. 4F). Co-treatment with both agonists together abolished the previously observed signal (Fig. 4F), indicating that NMDAR activation blunts CB_2_R-mediated signalling in MAPK pathway. Moreover, the signal observed upon JWH-133 treatment was blocked not only by CB_2_R antagonist SR144528, but also by NMDAR antagonist MK-801 (Fig. 4F), indicating that also in this signalling pathway there is a cross-antagonism effect from NMDAR to CB_2_R. When microglial cells were treated with α-syn fibrils, JWH-133 signal was diminished compared to control cells (Fig. 4G), suggesting that α-syn fibrils treatment decreases CB_2_R signalling through the MAPK pathway in microglial cells.

### 2.6. CB_2_R activation counteracts the detrimental phenotype of microglia activation induced by α-syn fibrils

Finally, we wondered the effect of α-syn fibrils on microglial polarization to either the detrimental M1 phenotype or the neuroprotective M2 phenotype, and if CB_2_R or NMDAR activation could counteract these effects. Therefore, primary cultures of microglia treated with α-syn fibrils and CB_2_R or NMDAR agonists were subjected to immunocytochemistry with antibodies for either Iba-1 as a general indicator of microglial activation, Inducible Nitric Oxide Synthase (iNOS) as a marker of M1 phenotype, or Arginase-1 (Arg-1) as a marker of M2 phenotype. When microglial cells were treated with α-syn fibrils, Iba-1 signal significantly increased, and this increase was not modified upon treatment with neither CB_2_R agonist JWH-133 nor with NMDA (Fig. 5A). Interestingly, treatment with NMDA alone also increased Iba-1 signal (Fig. 5A). As expected, α-syn fibrils also increased iNOS signal, and interestingly this effect was counteracted by CB_2_R but not by NMDAR agonism (Fig. 5B), which indicates that CB_2_R but not NMDAR activation can alleviate the polarization of microglia to the activated M1 phenotype. Finally, when analyzing Arg-1 signal, we could observe that it was slightly decreased by α-syn fibrils treatment (Fig. 5C). However, treatment with CB_2_R agonist JWH-133 significantly increased Arg-1 signal, effect that was not observed upon NMDA treatment (Fig. 5C), reinforcing the idea that cannabinoids have a neuroprotective effect by not only decreasing the polarization of microglia to M1 phenotype, but also by increasing the transition to M2 phenotype.

**Fig 5.**
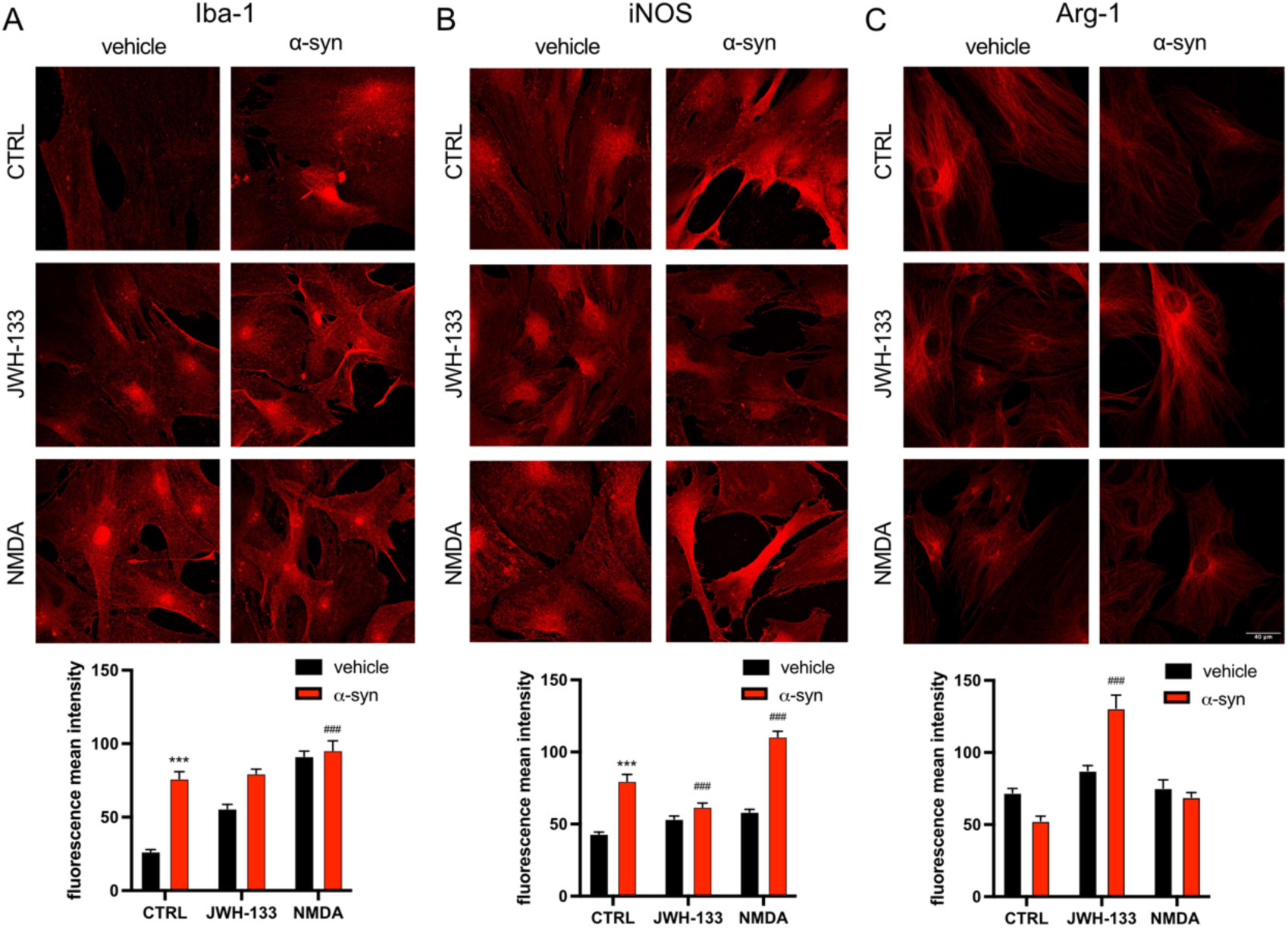
Analysis of the effects of α-syn fibrils on microglia polarization. Microglial markers in primary cultures treated with α-syn fibrils. Primary cultures of microglia were treated for 48 h with the specific agonist for CB_2_R, JWH-133 (100 nM) or for NMDAR, NMDA (15 mM) in the presence of α-syn fibrils. Immunocytochemical assays were performed using primary antibodies that detect either the Iba-1 microglial activation marker (A), the arginase-1 M1 marker (B) or the inducible nitric oxide synthase M2 marker (C) followed by Cy3 secondary antibody (red). Fluorescence was quantified using Fiji program (raw fluorescence measurements are shown). Representative images for all conditions are shown. Scale bar 40 μm. Two-way ANOVA followed by Tukey’s multiple comparison post hoc test were used for statistical analysis (***p < .001; versus vehicle, ###p < .001; versus control).

## 3. DISCUSSION

The interest in the use of cannabinoids as therapeutic agents is increasing, as new evidence appears showing the beneficial effects of these compounds in different pathologies (Maccarrone et al., 2023). Cannabinoids act mainly on cannabinoid receptors 1 and 2, which together with endocannabinoids and with the enzymes that synthetize and degrade these compounds form the endocannabinoid system. Parkinson’s disease courses with alterations in the endocannabinoid system. The levels of endocannabinoids are dysregulated in PD patients (A. Pisani et al., 2005; V. Pisani et al., 2010) and in rodents that overexpress α-synuclein (Kelly et al., 2022). CB_1_R expression changes along the progress of the pathology, being downregulated at the initial phase, and upregulated in the late stages, both in patients and animal models (Soti et al., 2022). Regarding CB_2_R, its expression has been found to be upregulated in microglia in the MPTP neurotoxic model of PD (Price et al., 2009), while other studies have also found an increase of CB_2_R expression in the striatum and substantia nigra of the inflammation-driven, rotenone and 6-OHDA animal models of the pathology (Concannon et al., 2015, 2016; C. García et al., 2011; Gómez-Gálvez et al., 2016). In patients of the disease, CB_2_R levels appear to be upregulated in microglia (Gómez-Gálvez et al., 2016) but downregulated in neurons of the substantia nigra (M. C. García et al., 2015).

These findings, together with the fact that NMDA receptors are also altered in Parkinson’s disease (Dunah et al., 2000; Gan et al., 2014; Mellone et al., 2015), led us to wonder if the complexes formed by CB_2_ and NMDA receptors could have a role in this pathology. When analyzing how the affinity of ligands for the orthosteric site of CB_2_R is affected in an in vitro PD model. α-synuclein preformed fibrils did not modify the affinity of the CB_2_R agonist JWH-133 for CB_2_R in HEK-293T cells expressing either CB_2_R alone or CB_2_R-NMDAR complexes. Hence, our observations showing that α-syn fibrils impair the signaling of CB_2_R, either when expressed alone or in combination with NMDAR, when analyzing intracellular cAMP accumulation, ERK1/2 phosphorylation and β-arrestin II recruitment in HEK-293T cells are not due to a decrease in the affinity of JWH-133 for CB_2_R. This reduction of cannabinoid receptor signalling upon α-syn fibrils incubation was previously detected for CB_1_R (Reyes-Resina et al., 2024).

When analyzing Ca^2+^ signaling, although increases Ca^2+^ intracellular levels have been described upon exposure of neurons to α-syn oligomers in striatal slices (Ronzitti et al., 2014), we observed a decrease of Ca^2+^ signaling in HEK-293T cells expressing CB_2_R-NMDAR complexes upon α-syn treatment. These results are intriguing but could be explained by the fact that HEK-293T cells lack many of the proteins present in neurons, thus the mechanisms controlling Ca^2+^ release in neurons are not present in HEK-293T cells. The most interesting observation when analyzing this pathway is the fact that activation of cannabinoid receptor 2 impairs NMDAR signal, as was described for CB_1_R-NMDAR complex (Reyes-Resina et al., 2024), and which highlight the dual benefits of cannabinoids: on one hand they provide the well know anti-inflammatory effects, and on the other hand they provide further benefit by decreasing NMDAR-induced excitotoxicity. Indeed, the use of NMDA receptor antagonist has proved to be effective for the treatment of PD symptoms (Armentero et al., 2006; Kelsey et al., 2004). There is evidence that NMDA receptor antagonism effectively improves motor symptoms in rodent PD models (Bortolanza et al., 2016; Egunlusi & Joubert, 2024; Flores et al., 2018; Konieczny et al., 1999). Selective NR2B antagonists, such as ifenprodil and traxoprodil, have shown efficacy in reducing L-DOPA-induced dyskinesia in both rodent models and MPTP-lesioned primates (Igarashi et al., 2015; Steece-Collier et al., 2000). Aslo, NR2B antagonists have shown promising results in combination therapy approaches. For instance, the co-administration of radiprodil with the A_2A_R antagonist tozadenant resulted in enhanced motor performance in 6-OHDA-lesioned rats (Michel et al., 2015).

α-syn fibrils affected not only receptor pharmacology, but they also induced a reorganization of CB_2_R-NMDAR complexes, as they increased the BRET signal in HEK-293T cells. This result apparently indicates that there is an increased heteromerization between CB_2_ and NMDAR induced by α-syn. However, PLA assays, that allow to detect heteromers in situ in native tissue, show that α-syn fibrils treatment significantly reduces the number of CB_2_R-NMDAR heteromers per cell in primary cultures of microglia, as previously observed for CB_1_R (Reyes-Resina et al., 2024). Hence, α-syn fibrils also affect heteromerization between CB_2_R and NMDAR.

Parkinson’s disease is known to course with neuroinflammation, as activated microglia has been found in the brains of PD patients (Gerhard et al., 2006; McGeer et al., 1988). Moreover, aggregated or mutated α-syn has been shown to induce microglial activation, which contributes to disease progression (Duffy et al., 2018; Grozdanov et al., 2019; Hoenen et al., 2016; Hoffmann et al., 2016; Zhang et al., 2005). In the CNS, the expression of CB_2_R is higher in glia than in neurons (Young & Denovan-Wright, 2022), being upregulated in microglia under inflammatory conditions in vivo (Concannon et al., 2015, 2016; Maresz et al., 2005; Medina-Vera et al., 2023; Solas et al., 2013) and in vitro (Navarro et al., 2018). While CB_2_R expression in the healthy brain is relatively low, both preclinical animal models of neurodegenerative diseases and brain tissue of PD patients present an elevated CB_2_R expression in microglia (Gómez-Gálvez et al., 2016). In line with these findings, we have observed that in microglia treated with α-syn fibrils, cAMP response to CB_2_R agonist is increased compared to control microglia, indicating that CB_2_R is able to elicit a stronger signal in the presence of α-syn fibrils in microglial cells. These results differ from those observed in HEK-293T cells, as microglial activation is strong enough to overcome the decrease in the affinity of CB_2_R agonists for the receptor. However, this stronger CB_2_R activation upon α-syn fibrils incubation was not observed when analyzing ERK1/2 phosphorylation in microglia.

Activated microglia can acquire a range of different phenotypes, being the proinflammatory M1 and the anti-inflammatory M2 opposed phenotypes. α-synuclein has been described to polarize activated microglia towards the detrimental M1 phenotype (Ardah et al., 2019; Codolo et al., 2013; Fellner et al., 2013; Kim et al., 2013; Reynolds et al., 2008; Rojanathammanee et al., 2011; Sarkar et al., 2020; Zhang et al., 2005), and M1 microglia has been found in the MPTP model of PD (Liberatore et al., 1999; McGeer & McGeer, 2008; Wu et al., 2003; Zhang et al., 2004). Furthermore, CSF and serum of PD patients show high levels of pro-inflammatory markers (Karpenko et al., 2018). On the contrary, microglia has been described to participate in the clearance of aggregated α-synuclein (Choi et al., 2020; George et al., 2019), therefore delaying the progression of the diseases. Specifically, α-synuclein clearance via M2 microglia polarization has been shown in experimental and human parkinsonian disorder (Park et al., 2016). When we analyzed microglial activation markers, we observed that α-syn fibrils indeed promoted microglial polarization towards M1 phenotype, effect that was counteracted by CB_2_R activation, but not by NMDAR activation. Moreover, activation of CB_2_R, increased polarization to the protective M2 phenotype, effect that was not observed upon NMDAR activation. These results point to a neuroprotective effect of CB_2_R activation in PD, as it was able to rescue the detrimental effects of α-syn fibrils in microglia. In line with these results, Mecha et al. demonstrated that CB_2_R expression in microglia is necessary to achieve M2 polarization (Mecha et al., 2015). Other studies have found increased expression of anti-inflammatory cytokines following CB_2_R specific activation (Correa et al., 2010; Ma et al., 2015; Malek et al., 2015).

Altogether, our results point to an anti-inflammatory and thus neuroprotective effect of CB_2_R activation that could delay the progression of the disease by counteracting the detrimental effects of oligomeric α-synuclein. Previous studies support the role of CB_2_R as therapeutic target for the treatment of PD, where CB_2_R activation resulted in reduced neuroinflammation, prevention of glial-derived neurotoxic mediator production, BBB leakage and peripheral immune cell infiltration, decreased death of dopaminergic neurons and alleviation of motor impairments in different animal models of PD (Chung et al., 2016; Gómez-Gálvez et al., 2016; Javed et al., 2016; Zhu et al., 2023). CB_2_R activation also promoted a reduction in the formation of alpha-synuclein aggregates in a rat model of nigral synucleinopathy (Joers et al., 2024). Moreover, the absence of CB_2_R impaired motor abilities, exacerbated the loss of dopaminergic neurons, and induced activation of the IL-1β pathway in the MPTP-induced mouse model of PD (Zhu et al., 2023). The lack of CB_2_R also resulted in an elevated α-synuclein-induced microglial synaptic pruning (Feng et al., 2023).

Nowadays there is no cure for PD, and no neuroprotective therapy to prevent neuronal loss is available. The most common strategy for the pharmacological treatment of PD tries to compensate for the loss of striatal dopamine by using the dopamine precursor levodopa (L-DOPA). However, the use of L-DOPA produces secondary undesired effects known as dyskinesias, consisting on abnormal involuntary movements (Ahlskog & Muenter, 2001; Bezard et al., 2001). The study by Rentsch et al. demonstrates that the CB_2_ agonist HU-308 was able to reduce levodopa-induced-dyskinesia in mice (Rentsch et al., 2020), which demonstrates that cannabinoids could not only address PD pathological hallmarks, but also those derived from the use of L-DOPA. As mentioned above, using NMDAR antagonist has shown benefits in PD treatment.

One of the current lines of research to find new PD therapies focuses on α-synuclein, both in immunotherapy with the use of antibodies directed against α-synuclein and in the use of inhibitors of α-synuclein aggregation. Some clinical trials have already reached phase II, but these drugs are not in the market yet (Calabresi et al., 2023). We propose the use of cannabinoids as new therapy for the treatment of PD. Cannabinoid action on CB_2_R could prevent neuronal death and promote polarization of microglia to the neuroprotective M2 phenotype. Indeed, both endocannabinoids, 2-arachidonoyl glycerol and anandamide, have antiparkinsonian properties; 2-AG is neuroprotective, and anandamide provides symptomatic relief (Abellanas & Aymerich, 2020).

## 4. MATERIALS AND METHODS

### 4.1. Drugs

JWH-133 (#1343), N-Methyl-D-aspartate (NMDA) (#0114), SR 144528 (#5039), (+)-MK 801 maleate (#0924) and zardaverine (#1046) were purchased from Tocris (Bristol, UK). Forskolin (FK) (#HY-15371/CS-1454) was purchased from MedChemExpress. The CB_2_R agonist 3-[[4-[2-tert-butyl-1-(tetrahydropyran-4-ylmethyl) benzimidazol-5-yl]sulfonyl-2-pyridyl]oxy]propan-1-amine (CM157) conjugated to a fluorescent probe was developed in collaboration with Cisbio Assays (Codolet, France) (see (Martínez-Pinilla et al., 2016)).

### 4.2. Cell Culture and Transient Transfection

HEK-293T cells at passage 8-12 were grown in Dulbecco’s modified Eagle’s medium (DMEM) (15-013-CV, Corning, NY, USA) supplemented with 2 mM L-glutamine, 100 U/ml penicillin/streptomycin, MEM Non-Essential Amino Acids Solution (1/100) and 5% (v/v) heat inactivated Fetal Bovine Serum (FBS) (Invitrogen, Paisley, Scotland, UK). Cells were maintained in a humid atmosphere of 5% CO2 at 37°C. Briefly, HEK-293T cells growing in 6-well dishes or in 25 cm2-flasks were transiently transfected using the PEI (PolyEthylenImine, Sigma-Aldrich, St. Louis, MO, USA) method. Cells were incubated for 4h with the corresponding cDNAs together with PEI (5.47 mM in nitrogen residues) and 150 mM NaCl in a serum-starved medium. Then, the medium was replaced by a fresh complete culture medium, and cells were incubated for 48 hrs before experimental procedures.

To prepare primary microglial cultures, brain was removed from 2-day-old Sprague Dawley rats. Brains were dissected, carefully stripped off the meninges and digested with 0.25% trypsin for 20 min at 37 °C. Trypsinization was stopped by washing the tissue. Cells were brought to a cell suspension by passage through 0.9 mm and 0.5 mm needles followed by passage through a 100 μm pore mesh. Glial cells were resuspended in medium and seeded at a density of 1 × 106 cells/ml in 6-well dishes for cyclic adenylic acid (cAMP) assays, in 12-well dishes with coverslips coated with poly-D-lysine (A38904-01, Gibco, Paisley, Scotland, UK) for in situ proximity ligation assays (PLA) and in 96-well plates for mitogen-activated protein kinase (MAPK) activation experiments. Cell counting was assessed using trypan blue and a Countess II FL automated cell counter (Thermo Fisher Scientific-Life Technologies). Cultures were grown in DMEM medium (15-013-CV, Corning, NY, USA) supplemented with 2 mM L-glutamine, 100 U/ ml penicillin/streptomycin, MEM non-essential amino acids preparation (1/100) and 5% (v/v) heat-inactivated fetal bovine serum (FBS) (Invitrogen, Paisley, Scotland, UK) and maintained at 37°C in humidified 5% CO2 atmosphere and, unless otherwise stated, medium was replaced once a week.

To prepare primary neuronal cultures, the brain from Sprague Dawley rat embryos (E19) was removed. The striatum was dissected and carefully stripped off the meninges. Tissue was processed as described above for microglial cultures, except that neurons were grown in a neurobasal medium (21103-049, Gibco, Paisley, Scotland, UK) supplemented with 2 mM L-glutamine, 100 U/ mL penicillin/streptomycin, MEM non-essential amino acids preparation (1/100), and 2% (v/v) B27 supplement (17504-044, Gibco, Paisley, Scotland, UK). Cultures were maintained at 37°C in a humidified 5% CO_2_ atmosphere for 12 days.

### 4.3. Fusion Proteins and Expression Vectors

The human cDNAs for the CB_2_ and GHSR_1A_ receptors and NR1A and NR2B NMDAR subunits cloned in pcDNA3.1 were amplified without their stop codons using sense and antisense primers. The primers harbored either unique BamHI and KpnI sites for CB_2_R, EcoRI and KpnI sites for GHSR_1A_ or BamHI and HindIII sites for NR1A. The fragments were subcloned to be in frame with an enhanced yellow fluorescent protein (pEYFP-N1; Clontech, Heidelberg, Germany) or an Rluc (pRluc-N1; PerkinElmer, Wellesley, MA) on the C-terminal end of the receptor to produce NR1A-Rluc, CB_2_R–YFP and GHSR_1A_-YFP fusion proteins. The plasmid encoding for the SNAP-tagged human CB_2_R used HTRF assays was obtained from Cisbio Bioassays (PSNAP-CB_2_, Cisbio Assays, Codolet, France).

### 4.4. α-synuclein fibrils treatment

HEK-293T or primary glial cell cultures were treated for 48h with recombinant human α-synuclein fibrils obtained by sonication, at a final concentration of 10 µg/L. Fibrils were prepared as previously described (Franco et al., 2018; Masuda-Suzukake et al., 2014).

### 4.5. Immunofluorescence Studies

HEK-293T or microglial cells growing on glass coverslips were fixed in 4% paraformaldehyde for 15 min and then washed twice with PBS containing 20 mM glycine before permeabilization with the same buffer containing 0.2% Triton X-100 (5 min incubation). Cells were treated for 1 h with PBS containing 1% bovine serum albumin. To detect the expression of NR1-Rluc, HEK-293T cells were labeled with a mouse anti-Rluc antibody (1/100; MAB4400, Millipore) and subsequently treated with Cy3-conjugated anti-mouse IgG secondary antibody (1/200; 715-166-150; Jackson ImmunoResearch) (1 h each). The expression of CB_2_R-YFP was detected by the YFP’s own fluorescence.

The presence of α-syn fibrils was detected with a mouse monoclonal anti-human α-synuclein antibody (1/300; ab1903, Abcam) followed by incubation with a Cy3-conjugated anti-mouse IgG secondary antibody (1/200; 715-166-150; Jackson ImmunoResearch) (1 h each). For M1/M2 assays, microglial cells were labeled with either a goat anti-Iba1 (1/100; ab107159; Abcam), a mouse anti-iNOS (1/100; NOS2 [C-11]: sc-7,271; SCB) or a mouse anti-arginase I (1/100; 610708; BD Biosciences) antibody, and subsequently treated with Cy3-conjugated anti-goat (1/200; 705-165-147; Jackson ImmunoResearch [red]) or anti-mouse (1/200; 715–166-150; Jackson ImmunoResearch [red]) IgG secondary antibodies (1 hr each).

Nuclei were stained with Hoechst 33432 (1/100 from stock 1 mg/mL; Thermo Fisher). The samples were washed several times and mounted on glass slides with ShandonTM Immu-MountTM (9990402; ThermoFisher). Samples were observed under a Zeiss 880 confocal microscope (Carl Zeiss, Oberkochen, Germany) equipped with an apochromatic 63× oil-immersion objective (N.A. 1.4), and with 405 nm, 488 nm and 561 nm laser lines.

### 4.6. Homogeneous competition binding assays

HEK-293T cells growing in 25 cm^2^ flasks were transiently transfected with either SNAP-CB_2_R or SNAP-CB_2_R together with NR1 plus NR2B subunits of NMDAR. For SNAP protein labeling, 48 h after transfection, cell culture medium was removed and 100 nM SNAP-Lumi4-Tb (SSNPTBC, Cisbio Assays, Codolet, France) diluted in Tag-lite Buffer (TLB, Cisbio Assays, Codolet, France) was added to the cells and incubated for 1 h at 37°C under 5% CO_2_ atmosphere. Cells were then washed with TLB to remove the excess of SNAP-Lumi4-Tb, detached with enzyme-free cell dissociation buffer, centrifuged 5 min at 1500 rpm and resuspended in TLB. To perform competition binding assays 2,500−3,000 cells/well were placed in white opaque 384-well plates. Then, 20 nM fluorophore-conjugated CB_2_R ligand (labeled CM157) diluted in TLB was added to the cells, followed by increasing concentrations (0–10 μM range) of unlabeled JWH-133. Plates were then incubated for 2 h at room temperature before signal detection. Homogeneous time-resolved fluorescence energy transfer (HTRF) was detected using a PHERAstar Flagship microplate reader (Perkin-Elmer, Waltham, MA, USA) equipped with a FRET optic module allowing donor excitation at 337 nm and signal collection at both 665 and 620 nm.

### 4.7. Bioluminescence Resonance Energy Transfer (BRET) Assays

HEK-293T cells growing in 6-well plates were transiently co-transfected with a con-stant amount of cDNA encoding for NR1A fused to Renilla luciferase (NR1A-Rluc), with a constant amount of the cDNA encoding for NR2B, and with increasing amounts of cDNAs corresponding to CB_2_R or ghrelin receptor GHSR_1A_ fused to the yellow fluorescent protein (CB_2_-YFP, GHSR_1A_-YFP). 48 h post-transfection cells were washed twice in quick succession in HBSS (137 mM NaCl; 5 mM KCl; 0.34 mM Na2HPO4; 0.44 mM KH2PO4; 1.26 mM CaCl2; 0.4 mM MgSO4; 0.5 mM MgCl2 and 10 mM HEPES, pH 7.4) supplemented with 0.1% glucose (w/v), detached by gently pipetting and resuspended in the same buffer. To assess the number of cells per plate, we determined protein concentration using a Bradford assay kit (Bio-Rad, Munich, Germany) with bovine serum albumin dilutions as standards. To quantify YFP-fluorescence expression, we distributed the cells (20 μg protein) in 96-well microplates (black plates with a transparent bottom; Porvair, Leatherhead, UK). Fluorescence was read using a Mithras LB 940 (Berthold, Bad Wildbad, Germany) equipped with a high-energy xenon flash lamp, using a 10-nm bandwidth excitation and emission filters at 485 and 530 nm, respectively. YFP-fluorescence ex-pression was determined as the fluorescence of the sample minus the fluorescence of cells expressing protein-Rluc alone. For the BRET measurements, the equivalent of 20 μg of cell suspension was distributed in 96-well microplates (white plates; Porvair), and we added 5 μM coelenterazine H (PJK GMBH, Kleinblittersdorf, Germany). Then, 1 min after coelenterazine H addition, the readings were collected using a Mithras LB 940 (Berthold, Bad Wildbad, Germany), which allowed the integration of the signals detected in the short-wavelength filter at 485 nm (440–500 nm) and the long-wavelength filter at 530 nm (510–590 nm). To quantify receptor-Rluc expression, we performed luminescence read-ings 10 min after addition of 5 μM coelenterazine H. The net BRET is defined as [(long-wavelength emission)/ (short-wavelength emission)]-Cf where Cf corresponds to [(long-wavelength emission)/ (short-wavelength emission)] for the Rluc construct ex-pressed alone in the same experiment. The BRET curves were fitted assuming a single phase by a non-linear regression equation using the GraphPad Prism software (San Diego, CA, USA). BRET values are given as milli BRET units (mBU: 1000 × net BRET).

### 4.8. cAMP Level Determination

Two hours before initiating the experiment, HEK-293T or microglial cell-culture medium was exchanged to serum-starved DMEM or neurobasal medium, as corresponds. Then, cells were detached, resuspended in the serum-starved medium containing 50 µM zardaverine and plated in 384-well microplates (2,500 cells/well), pretreated (15 min) with the corresponding antagonists— or vehicle—and stimulated with agonists (15 min) before adding 0.5 μM forskolin or vehicle. Readings were performed after 1 h incubation at 25 °C. Homogeneous time-resolved fluorescence energy transfer (HTRF) measures were obtained using the Lance Ultra cAMP kit (PerkinElmer, Waltham, MA, USA) [62]. Fluorescence at 665 nm was analyzed on a PHERAstar Flagship microplate reader equipped with an HTRF optical module (BMG Lab technologies, Offenburg, Germany).

### 4.9. Extracellular Signal-Regulated Kinase Phosphorylation Assays

HEK293T cells growing in 25 cm^2^ flasks were transfected with the cDNAs encoding for CB_2_R, for NR1A and for NR2B. Two to four hours before initiating the experiment, the culture medium was replaced by serum-starved DMEM medium. Cells were incubated at 37°C with antagonists (15 min) or vehicle, followed by stimulation (7 min) with agonists. After that, the reaction was stopped by placing cells on ice. Then, cells were washed twice with cold PBS and lysed by the addition of ice-cold lysis buffer (50 mM Tris-HCl pH 7.4, 50 mM NaF, 150 mM NaCl, 45 mM glycerol-3-phosphate, 1% Triton X-100, 20 µM phenyl-arsine oxide, 0.4 mM NaVO4 and protease inhibitor mixture (MERK, St. Louis, MO, USA)) Cellular debris were removed by centrifugation at 13,000 g for 10 min at 4 °C, and protein concentration was adjusted to 1 mg/mL by the bicinchoninic acid method (ThermoFisher Scientific, Waltham, MA, USA) using a commercial bovine serum albumin dilution (BSA) (ThermoFisher Scientific) for standardization. 6x Laemli SDS sample buffer (300 mM Tris-Base, 600 mM DTT, 40% glycerol (v/v), 0,012 % Bromophenol blue (w/v) and 12 % SDS (w/v), pH= 6,8) was added to the samples and proteins were denatured by boiling at 100 °C for 5 min. ERK1/2 phosphorylation was determined by Western blot. Equivalent amounts of protein (20 μg) were subjected to electrophoresis (10% SDS-polyacrylamide gel) and transferred onto PVDF membranes (Immobilon-FL PVDF membrane, MERK, St. Louis, MO, USA) for 30 min using Trans-Blot Turbo system (Bio-Rad). Then, the membranes were blocked for 2 h at room temperature (constant shaking) with Odyssey Blocking Buffer (LI-COR Biosciences, Lincoln, NE, USA) and labeled with a mix of primary mouse anti-phospho-ERK 1/2 (1/2500, MERK, Ref. M8159) and rabbit anti-ERK 1/2 (1/40,000, MERK, Ref. M5670) antibodies overnight at 4 °C with shaking. Then, the membranes were washed three times with PBS containing 0.05% tween for 10 min and subsequently were incubated with a mix of IRDye 800 anti-mouse (1/10,000, MERK, Ref. 92632210) and IRDye 680 anti-rabbit (1/10,000, MERK, Ref. 926-68071) secondary antibodies for 2 h at room temperature, light protected. Membranes were washed 3 times with PBS-tween 0.05% for 10 minutes and once with PBS and left to dry. Bands were analyzed using Odyssey infrared scanner (LI-COR Biosciences). Band densities were quantified using Fiji software, and the level of phosphorylated ERK1/2 was normalized using the total ERK 1/2 protein band intensities. Results are represented as the percent over basal (non-stimulated cells).

To determine extracellular signal-regulated kinase 1/2 (ERK1/2) phosphorylation in microglial primary cultures, cells were grown in 96-well plates. On the day of the experiment the medium was replaced by serum-free medium 2h before starting the experiment. Cells were pre-treated at 25 °C for 15 min with antagonists or vehicle and stimulated for an additional 15 min with selective agonists. Cells were then washed twice with cold PBS before the addition of lysis buffer (a 15 min treatment). Afterward, 10 µL of each supernatant were placed in white ProxiPlate 384-well plates and ERK 1/2 phosphorylation was determined using an AlphaScreen®SureFire® kit (Perkin Elmer), following the instructions of the supplier, and readings were collected using an EnSpire® Multimode Plate Reader (PerkinElmer, Waltham, MA, USA). The value of reference (100%) was the value achieved in the absence of any treatment (basal). The effect of ligands was given in percentage with respect to the basal value.

### 4.10. Detection of cytoplasmic calcium levels

HEK-293T cells were cotransfected with the cDNA for the corresponding receptors (see figure legend) together with the cDNA for the GCaMP6 calcium sensor (T.-W. Chen et al., 2013). 48 hours after transfection, HEK-293T cells were detached using Mg^2+^-free Locke’s buffer (154 mM NaCl, 5.6 mM KCl, 3.6 mM NaHCO3, 2.3 mM CaCl2, 5.6 mM glucose, 5 mM HEPES, 10 μM glycine, pH 7.4), centrifuged for 5 min at 3,200 rpm and resuspended in the same buffer. Protein concentration was quantified by using the Bradford assay kit (Bio-Rad, Munich, Germany). To measure Ca^2+^ mobilization, cells (40 µg of protein) were distributed in 96-well microplates (black plates with a transparent bottom; Porvair, Leatherhead, UK) and were incubated for 10 min with antagonists when indicated. Fluorescence readings were performed right after the addition of agonists. Fluorescence emission intensity due to GCaMP6 was recorded at 515 nm upon excitation at 488 nm on the EnSpire® Multimode Plate Reader for 300 s every 5 s at 100 flashes per well.

### 4.11. β-arrestin II recruitment

To determine β-arrestin recruitment, BRET experiments were performed in HEK-293T cells 48 h after transfection with the cDNA for either CB_2_R-YFP, or CB_2_R-YFP and NR1 and NR2B subunits of NMDAR, together with cDNA corresponding to β-arrestin II-RLuc. Cells (20 μg protein) were distributed in 96-well microplates (Corning 3600, white plates with white bottom) and incubated with antagonists for 10 min prior to the addition agonists. Inmediately after, 5 μM coelenterazine H (Molecular Probes, Eugene, OR) was added, and 7 min after coelenterazine H addition, BRET readings corresponding to β-arrestin II-Rluc and receptor-YFP were quantified. The readings were collected using a Mithras LB 940 (Berthold Technologies, Bad Wildbad, Germany) that allows the integration of the signals detected in the short-wavelength filter at 485 nm and the longwavelength filter at 530 nm. To quantify protein-RLuc expression, luminescence readings were performed 10 min after addition of 5 μM coelenterazine H.

### 4.12. Cell viability

Cell viability assay is based on the principle that living cells maintain intact cell membranes that exclude certain dyes, like trypan blue. To quantify the percentage of living cells, DIV 14 primary cultures of striatal neurons growing in 6-well plates were treated with 10 µg/L α-syn or vehicle On DIV 15, cells were treated either with 100 nM ACEA, 15 µM NMDA or vehicle for another 24 hrs.

On the day of the experiment, cells were gently detached and mixed with an equal volume of trypan blue (0.4%) (Trypan Blue solution, T8154, Sigma Aldrich (St. Louis, MO, US)). Cells (%) were counted in a Countess II FL automated cell counter (Thermo Fisher Scientific).

### 4.13. In Situ and In Vitro Proximity Ligation Assay (PLA)

The proximity ligation assay (PLA) allows the detection of molecular interactions between two endogenous proteins ex vivo. PLA requires both receptors to be sufficiently close (<16 nm) to allow double-strand formation of the complementary DNA probes conjugated to the antibodies. Using the PLA, the heteromerization of NR1 subunits of NMDAR with CB_2_ receptors was detected in situ in primary cultures of neurons and in rat brain sections.

The presence/absence of receptor–receptor molecular interactions in the samples was detected using the Duolink II In Situ PLA Detection Kit (developed by Olink Bioscience, Uppsala, Sweden; and now distributed by Sigma-Aldrich as Duolink® using PLA® Technology). The PLA probes were obtained after conjugation of the primary anti-NR1 antibody (ab52177, Abcam, Cambridge, UK) to a MINUS oligonucleotide (DUO92010, Sigma-Aldrich, St. Louis, MO, USA), and the anti-CB_2_R (ab230791, Abcam, Cambridge, UK) antibodies to a PLUS oligonucleotide (DUO92009, Sigma-Aldrich, St. Louis, MO, USA). The specificity of antibodies was tested in non-transfected HEK-293T cells. Microglial cells growing in glass coverslips were fixed in 4% paraformaldehyde for 15 min and then washed twice with PBS containing 20 mM glycine before permeabilization with PBS-glycine containing 0.2% Triton X-100 for 5 min. After permeabilization, samples were washed in PBS at room temperature and incubated in a preheated humidity chamber for 1 h at 37 °C with the blocking solution provided in the PLA kit. Then, samples were incubated overnight with the PLA probe-linked antibodies (1/100 dilution for all antibodies) at 4 °C. After washing, samples were incubated with the ligation solution for 1 h, and then washed and subsequently incubated with the amplification solution for 100 min (both steps at 37 °C in a humid chamber). Nuclei were stained with Hoechst33432 (1/100 from stock 1 mg/mL; Thermo Fisher). The samples were washed several times and mounted on glass slides with ShandonTM Immu-MountTM (9990402; ThermoFisher). Negative controls were performed by omitting the anti-CB_2_R-PLUS antibody. Samples were observed under a Zeiss 880 confocal microscope (Carl Zeiss, Oberkochen, Germany) equipped with an apochromatic 63× oil-immersion objective (N.A. 1.4), and with 405 and 561 nm laser lines. For each field of view, a stack of two channels (one per staining) and 9 Z planes with a step size of 0.5 µm were acquired. The ratio r (number of red spots/cell) was determined on the maximum projection of each image stack using the Duolink Image tool software.

### 4.14. Data Analysis

Data, expressed as the mean ± SEM, were obtained from at least five independent experiments. Data comparisons were analyzed by one-way ANOVA or two-way ANOVA, followed by Bonferroni’s post-hoc test. The normality of populations and homogeneity of variances were tested before the ANOVA. Statistical analysis was under-taken only when each group size was at least n = 5, n being the number of independent variables (technical replicates were not treated as independent variables). Differences were considered significant when P ≤ 0.05. Statistical analyses were carried out with GraphPad Prism software version 5 (San Diego, CA, USA). Outliers’ tests were not used, and all data points (mean of replicates) were used for the analyses.

## Author Contributions

Conceptualization, I.R-R. and G.N.; methodology, I.R-R. and G.N.; validation, I.R-R. and G.N.; formal analysis, I.R-R., J.L., and G.N.; investigation, J.L., J.B.R., I.R., T.C., P.B. and I.R.-R.,; data curation, J.L., J.B.R., I.R., T.C., P.B. and I.R.-R.; writing—original draft preparation, I.R-R. and G.N.; writing—review and editing J.L., J.B.R., I.R., T.C., P.B. and I.R.-R..; visualization, J.L., I.R. and J.B.R.; supervision, I.R-R. and G.N.; project administration, I.R-R. and G.N.; funding acquisition G.N. All authors have read and agreed to the published version of the manuscript.

## Data Availability Statement

Data can be obtained from the corresponding author upon reasona-ble request.

